# The ENIGMA-PD-WML Pipeline: A Containerized, User-Friendly Approach for Accurate, Standardized Segmentation of White Matter Lesions in Multi-Site MRI Data

**DOI:** 10.64898/2026.06.11.731538

**Authors:** Sarah Al-Bachari, Shauna Angell, Aswin Abraham, Yahya Khubrani, Paul Smith, Kimberly Meechan, Robin Long, Sunanda Somu, Renee Mapa, Conor Owens-Walton, Elizabeth Haddad, Sophia I. Thomopoulos, Carole H. Sudre, Ludovica Griffanti, Hosung Kim, Gilsoon Park, Ysbrand D. van der Werf, Paul M. Thompson, Neda Jahanshad, Chris Vriend, Anette Schrag, Hamied A. Haroon

## Abstract

Understanding vascular contributions to disease is a major unmet need. White matter lesions (WML) are an accepted imaging marker of cerebral small vessel disease, giving insights into its related pathologies. A unified approach for WML analyses in large multi-site data is lacking despite the need for pooling of data to overcome the limitations of often small heterogenous MRI studies which make subtyping and identifying patterns within disease groups difficult.

Our ENIGMA-PD-WML pipeline is an open-source containerized pipeline containing all the code and packages required for pre-processing, processing and post-processing of T1-weighted and FLAIR data, outputting accurate and reproducible binary WML maps using a UNet approach. The pipeline provides a standardized image analysis approach for WML and outputs data in both native and MNI space to allow for sharing and pooling of data from multiple sites for large-data analysis.

In addition to a reliable standardized approach for WML segmentation, key priorities when developing the pipeline included: usability, i.e., requiring minimal manual input and technical expertise to use, and suitability to run on various MRI scanners and acquisition parameters as is common in multi-site data. This paper describes the pipeline in detail, with rationale for each step, providing transparency and facilitating its usage to overcome reproducibility issues in large-scale WML analyses.

## Introduction

White matter lesions (WML) appear as hyperintensities on T2-weighted fluid-attenuated inversion recovery (FLAIR) and hypointensities on T1-weighted magnetic resonance imaging (MRI). Due to the increased water content of WML, they are most readily visible on FLAIR sequences, in which the signal from the cerebrospinal fluid is attenuated; however, combining FLAIR and T1-weighted images can enhance the contrast and anatomical detail, reducing the risk of misidentifying WML and ensuring the observed hyperintensities are not artefacts (Li et al., 2018). Traditionally WML have received most attention in neuroinflammatory disorders such as multiple sclerosis (MS). However, more recently, WML of presumed vascular origin have become an accepted surrogate marker of small vessel disease and have received much interest, especially within neurodegenerative disorders such as Alzheimer’s and Parkinson’s disease, cognitive syndromes, and in association with vascular risk factors and are therefore the focus of this pipeline (Shi and Wardlaw, 2016, Wardlaw et al., 2013, Garnier-Crussard et al., 2022, Reheman et al., 2025, Scamarcia et al., 2021, Lee and Lee, 2016). WML measurement is non-invasive and quantifiable with MR protocols commonly used in research and in the clinical setting, making WML load an ideal imaging marker of vascular health in human studies and case-control analyses. Of note, WML of presumed vascular origin often have distinguishable imaging appearances, patterns and progression from neuroinflammatory causes (Zhang et al., 2024).

Accumulating evidence suggests an association between the presence of WML of presumed vascular origin and various clinically important outcomes including cognitive decline, dementia, depression, and motor dysfunction (d’Arbeloff et al., 2019, Firbank et al., 2005, Gu et al., 2025, Hu et al., 2021, Lee et al., 2020, Roseborough et al., 2023). These findings suggest that WML can act as markers to assess how vascular dysfunction impacts important neurological functions. Lesion location, whether periventricular or in deep white matter, has been linked to differing pathologies (Kim et al., 2008, Cai et al., 2022).

Despite the growing interest in WML and their association with disease etiology, progression and clinical features, a standardized analysis approach is lacking. The current gold standard is manual segmentation, in which a trained radiologist manually delineates each lesion on a scan. However, this process is time and labor intensive, requires significant training and expertise, and is susceptible to inter- and intra-rater variability (Caligiuri et al., 2015). Consequently, there is a need for an automated segmentation approach to enable the assessment of WML in large, multi-site datasets.

The importance of large-data analysis is increasingly recognized. Most MRI studies are small, making it difficult to subtype and identify disease patterns in heterogeneous populations (Shawa et al., 2025). Our previous work focused on comparing various freely available automated WML approaches specifically in multi-site data with heterogeneous scanners and image acquisition parameters (Al-Bachari et al., 2026b). Recent studies have revealed the importance of not only overall lesion burden but also location, especially in strategic pathways, therefore both volumetric and spatial agreement with the gold standard of manual segmentation were assessed in our study (Lemprière, 2023, Melazzini et al., 2021). We used open-source datasets, comprising a Parkinson’s disease and healthy ageing population and found that a UNet approach outperformed other methods on various performance metrics, including Dice score, Hausdorff distance, intraclass correlation coefficient, F1, precision and recall (Al-Bachari et al., 2026b). The UNet approach produced accurate and reliable WML maps despite being subjected to various scanner parameters and lesion loads. Our findings are largely in keeping with other assessments of automated approaches (Kuijf et al., 2019, Kuijf, 2022).

UNet is a type of convolutional neural network, a deep learning image segmentation method that has repeatedly performed well for WML segmentations in various disease cohorts (Ronneberger et al., 2015). Li et al. (2018) used an ensemble approach to improve the accuracy of UNet predictions. Li et al.’s ensemble UNet approach has been tested on 3D T1-weighted and 2D multi-slice FLAIR images of varying voxel sizes from 170 subjects obtained from 5 MR scanners of varying field strength at the WMH Segmentation Challenge 2017 (Kuijf et al., 2019), outperforming all other methods. UNet-pgs is an adapted version of Li et al.’s ensemble approach, trained using a highlighting foreground method to minimize partial volume effects at lesion boundaries (Park et al., 2021). Importantly, UNet-pgs has been validated across multiple disease cohorts including normal aging, mild cognitive impairment and Alzheimer’s disease from both the WMH Segmentation Challenge dataset, and an additional 243 subjects from the Alzheimer’s Disease Neuroimaging Initiative database (ADNI). Of note, various adapted versions of UNet have been used in MS (Belwal and Singh, 2025, Wahlig et al., 2023) but UNet-pgs focuses on WML of presumed vascular origin.

The Enhancing NeuroImaging Genetics through Meta-Analysis (ENIGMA) consortium is a collaborative network that brings clinical and pre-clinical researchers in various disease groups together (https://enigma.ini.usc.edu/). ENIGMA-PD has produced the largest MR dataset in Parkinson’s involving over 40 cohorts in over 20 countries (https://enigma.ini.usc.edu/ongoing/enigma-parkinsons/). To take advantage of multi-site data, there has been a move towards standardizing image analysis approaches by creating containerized imaging pipelines (Waller et al., 2022, Hu et al., 2023, Ren et al., 2025). A pipeline takes the raw images, applies pre-processing, image analysis and post-processing steps then generates results. This avoids the need for individual groups to develop and perform complex image analysis and allows for integration of comparable data, despite diversity and heterogeneity in disease groups and scanner parameters. Applying these pipelines to multi-site data can standardize the processing, and has yielded important results, highlighting their value in identifying disease-specific patterns (Kerestes et al., 2023, Owens-Walton et al., 2024, Laansma et al., 2024). A containerized approach contains all of the software packages required, ensuring processing is identical across different research groups, allowing version control and producing consistent and reproducible outputs (Matelsky et al., 2018). Following the successful examples of fully containerized image analysis pipelines created by various groups within ENIGMA, we developed the ENIGMA-PD-WML pipeline.

The ENIGMA-PD-WML pipeline facilitates the use of FSL (version 6.0.7.22) tools (Jenkinson et al., 2012) alongside the UNet-pgs algorithm (Park et al., 2021) to allow automated segmentation and generation of binary WML maps from an individual subject’s T1-weighted and FLAIR images. The only minimum imaging requirements for the ENIGMA-PD-WML pipeline are a field strength of 1.5T or greater and a slice thickness of 5 mm or less. The final binary WML maps are output in MNI standard space allowing non-identifiable derived imaging data to be shared between sites minimizing the ethical, legal and financial constraints required for raw, identifiable data.

Having previously found UNet-pgs to outperform other automated WML segmentation approaches in terms of accuracy regardless of lesion load or location (Al-Bachari et al., 2026b), our key priorities for the development of the ENIGMA-PD-WML segmentation pipeline were:

1. A user-friendly approach, meaning any site, regardless of level of technical expertise, can use the containerized pipeline easily, with minimal manual input.
2. A pipeline that is robust against a range of MRI parameters and scanners as is reflective of multi-site data.

The following sections outline the practical steps to setting up the pipeline, followed by an in-depth explanation of the pre- and post-processing steps. For each step, we provide the rationale alongside detailed diagrams, aiming for glass-box transparency to encourage the use of the pipeline by fellow researchers and clinicians alike.

### Preparing and Running the Pipeline

#### Pipeline Prerequisites

For input into the pipeline, whole-brain T1-weighted and T2-weighted FLAIR (FLAIR) images must be in Neuroimaging Informatics Technology Initiative (NIfTI) format and organized in a way which allows the pipeline to automatically read the images. To increase the usability and reach, the pipeline has been developed to allow various data structures. There has been a recent move towards standardizing neuroimaging data organization, recommending data are structured according to BIDS format (Gorgolewski et al., 2016) as shown in **Figure 1**. Data for the pipeline can be organized in a simple BIDS format which does not require accompanying meta-data files.

**Figure 1:**
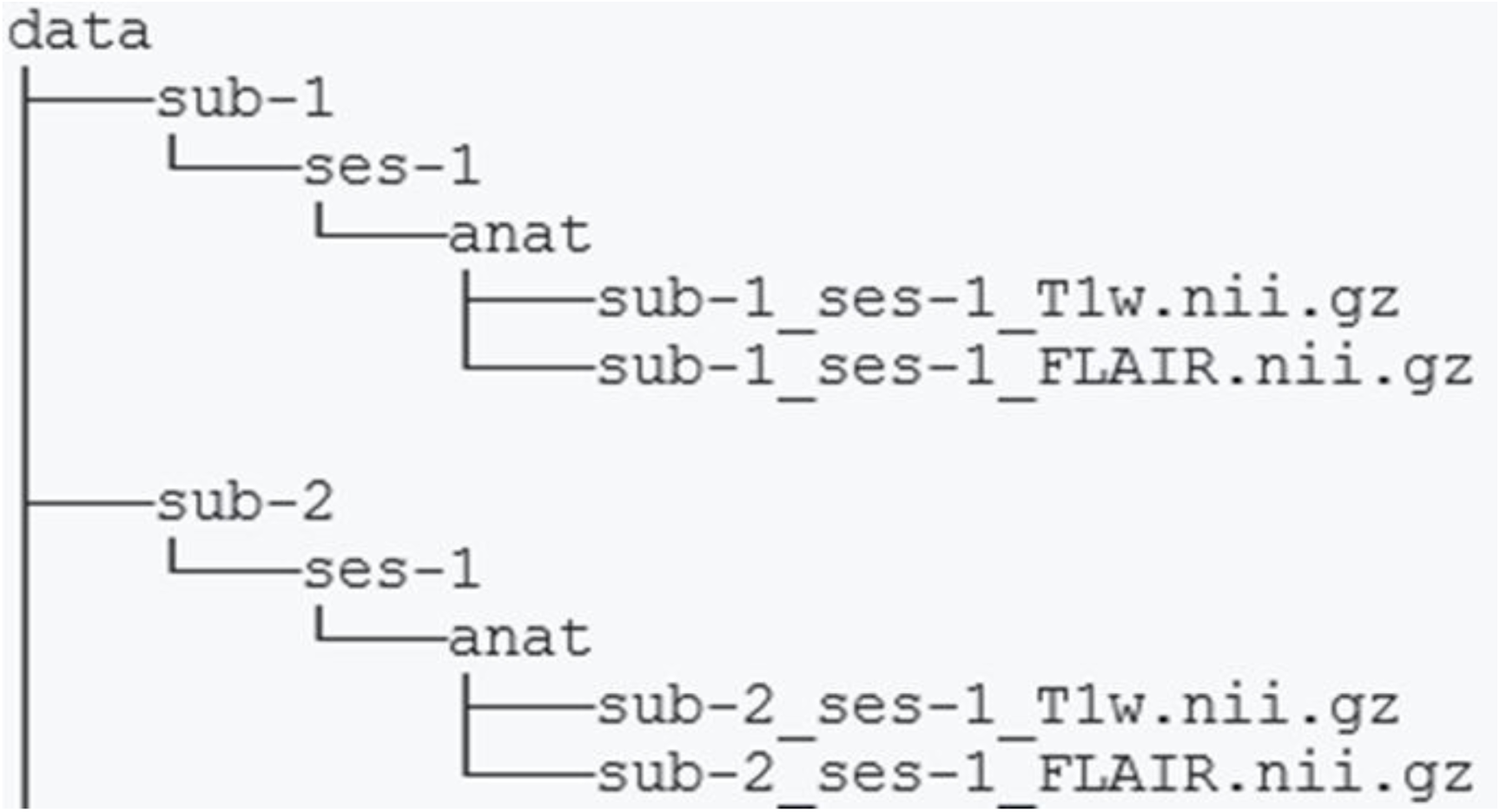
A representation of data organized in BIDS format for use in the ENIGMA-PD-WML pipeline.

Reorganization of data is the most manually intensive process of the pipeline. To allow flexibility in data structure, the pipeline accepts data in formats other than BIDS, thereby minimizing efforts from individual sites who may have their data stored in a site-specific manner. This requires creation of a CSV file that describes the directory structure of the data. Detailed instructions on this process are available on GitHub (https://github.com/UCL/Enigma-PD-WML/blob/main/docs/non-bids-data.md).

#### Implementing the Pipeline

The pipeline itself and instructions for use are freely available on GitHub (https://github.com/UCL/Enigma-PD-WML). The pipeline is available as both a Docker and Apptainer (formerly Singularity) container, both of which are free to use, allowing for wider usage on many different systems (https://www.docker.com/, https://apptainer.org/). Neither environment is superior, and use can be decided based on availability, preference and experience. Often Apptainer is used on high performance computing (HPC) systems because of root (admin) access restrictions for Docker. Regardless of the environment used, the pipeline must be run from the top-level directory, typically the “data” folder as described above.

#### Docker

If using Docker, freely available Docker application and installation instructions are available for Mac, Windows and Linux operating systems (https://docs.docker.com/get-started/get-docker/). Once installed, the pipeline image can be accessed from the enigma-pd-wml repository on Docker Hub (https://hub.docker.com/r/hamiedaharoon24/enigma-pd-wml/tags). It is recommended to use the latest version of the Docker image, which sits at the top of the repository page. The image can then be copied and run from the command line. An example of the command line is shown below:

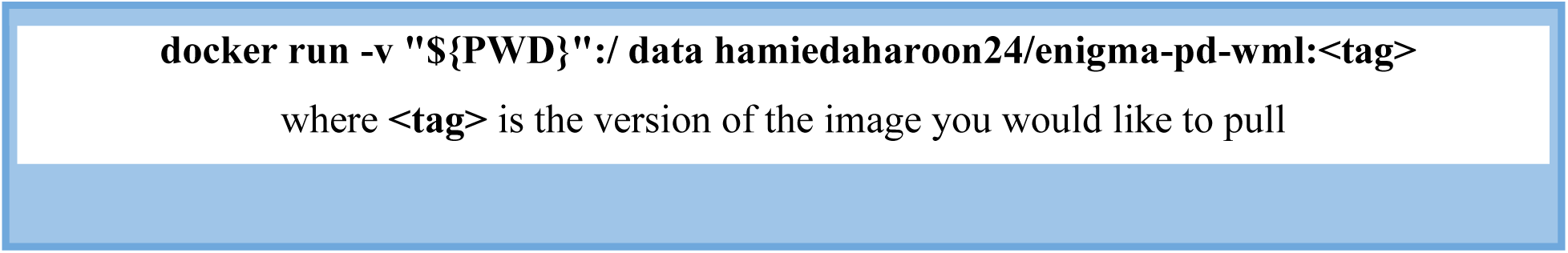

The first time this command is run the image will be downloaded. This may take some time, following which the pipeline will automatically begin running through the entire process.

#### Apptainer (Formerly Singularity)

Alternatively, if the user prefers Apptainer, a sif image must be built using the image from Docker Hub, using the command below:

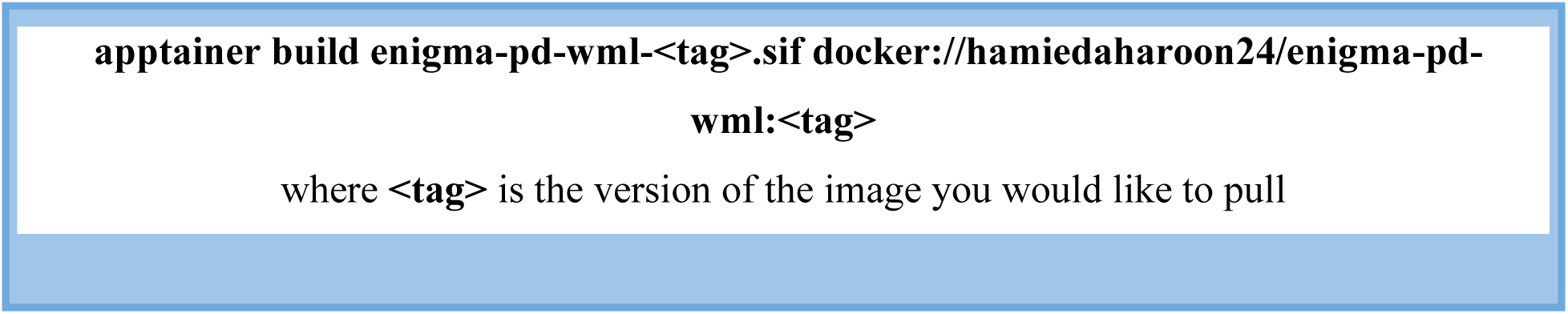

Similarly to Docker, using the latest version is recommended. This command will create an enigma-pd-wml-<tag>.sif file in the current working directory, which should be the top-level directory. Following this, the pipeline itself can be run using the following command:

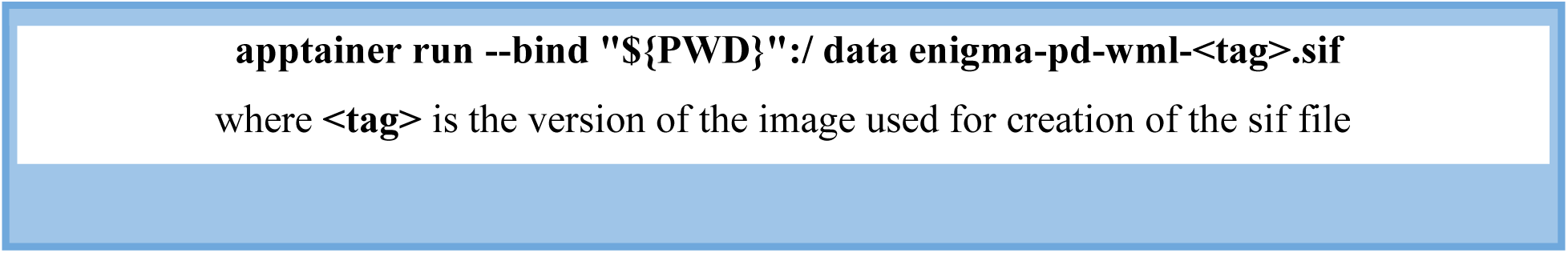

*Tip: The ENIGMA-PD-WML pipeline will be updated regularly with ongoing improvements, so it is important to ensure all subjects are run using the same version of the pipeline. Users should keep a note of the version used, in case any subjects need to be re-run or any issues arise at a later stage. All previous versions of the pipeline can be accessed on the GitHub page and Docker Hub. The details in this paper refer to version 1.1.1*.

Regardless of environment option, both of these processes will allow the pipeline to run on all the subjects for whom there are T1-weighted and FLAIR images present in the directory. **Table 1** describes the various command line options available to alter the default running of the pipeline.

**Table 1:** List of optional command line flags which can be included when running the ENIGMA-PD-WML pipeline and descriptions of their use.

| Optional Flag | Description |
| --- | --- |
| -n | Indicates the number of jobs the user wishes to run in parallel, defaulting to 1. Running the pipeline in parallel will shorten the overall time taken but also requires multiple cores and greater memory usage. |
| -o | Runs all steps of the pipeline, overwriting any previous steps. The default is for the pipeline to skip previously completed steps and re-use existing output files to reduce the time taken to re-run the pipeline if it fails at a late stage. |
| -f | Includes a path to a .txt file containing a list of subjects the pipeline is to be run on. Useful when the user only wants to analyze a subset of subjects in the directory. |
| -s | Includes a list of subjects on the command line for the pipeline to be run on. Similar to -f, this allows a subset of subjects to be analyzed. Each subject name in the list must be separated by a comma. |
| -l | This is required when using data in a non-BIDS format to allow the pipeline to recognize the data in the subject CSV file. |
| -h | This is required to provide a prefix for the HTML files generated in the quality control workflow when running the pipeline on different datasets. |

As the pipeline is fully containerized, the above describes all the manual input required in order to prepare the data and run the pipeline. Below we explain the processes within the pipeline which are automatically undertaken when the container is running.

A rationale throughout is that all outputs are produced in T1, FLAIR and MNI spaces, to enable quality control, sharing of de-identified data and maximum flexibility in subsequent analyses.

### Pre-processing, WML Segmentation and Post-processing

#### Pre-processing

**Figures 2, 3** and **4** provide a schematic overview of the preprocessing steps embedded in the pipeline for the T1-weighted images, FLAIR images and creation of WM masks respectively.

**Figure 2:**
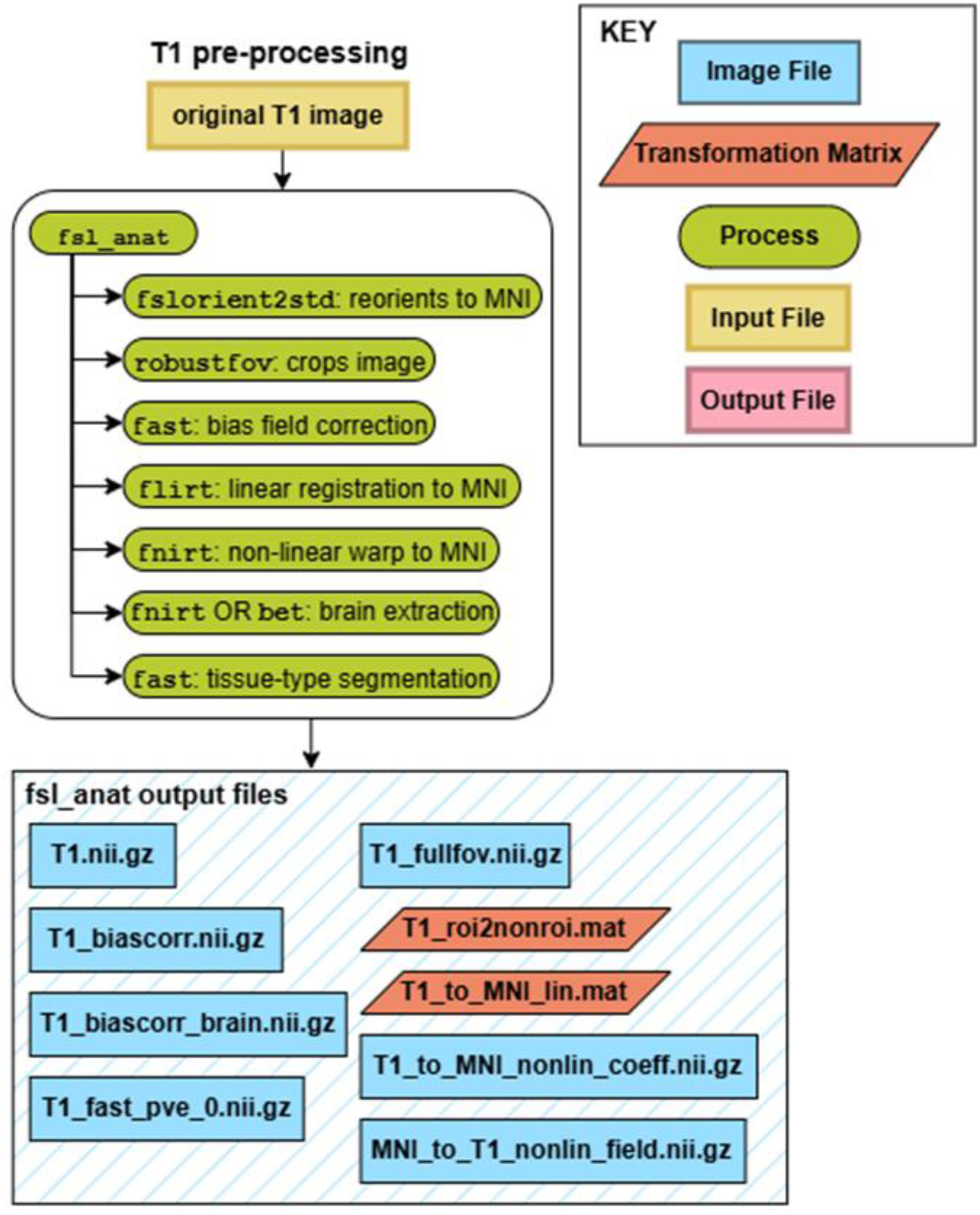
Schematic overview of the preprocessing of the T1-weighted images in the ENIGMA-PD-WML pipeline. Blue boxes show image files, red parallelograms show transformation matrices, green ovals show processes, yellow boxes show input files and pink boxes show output files.

**Figure 3:**
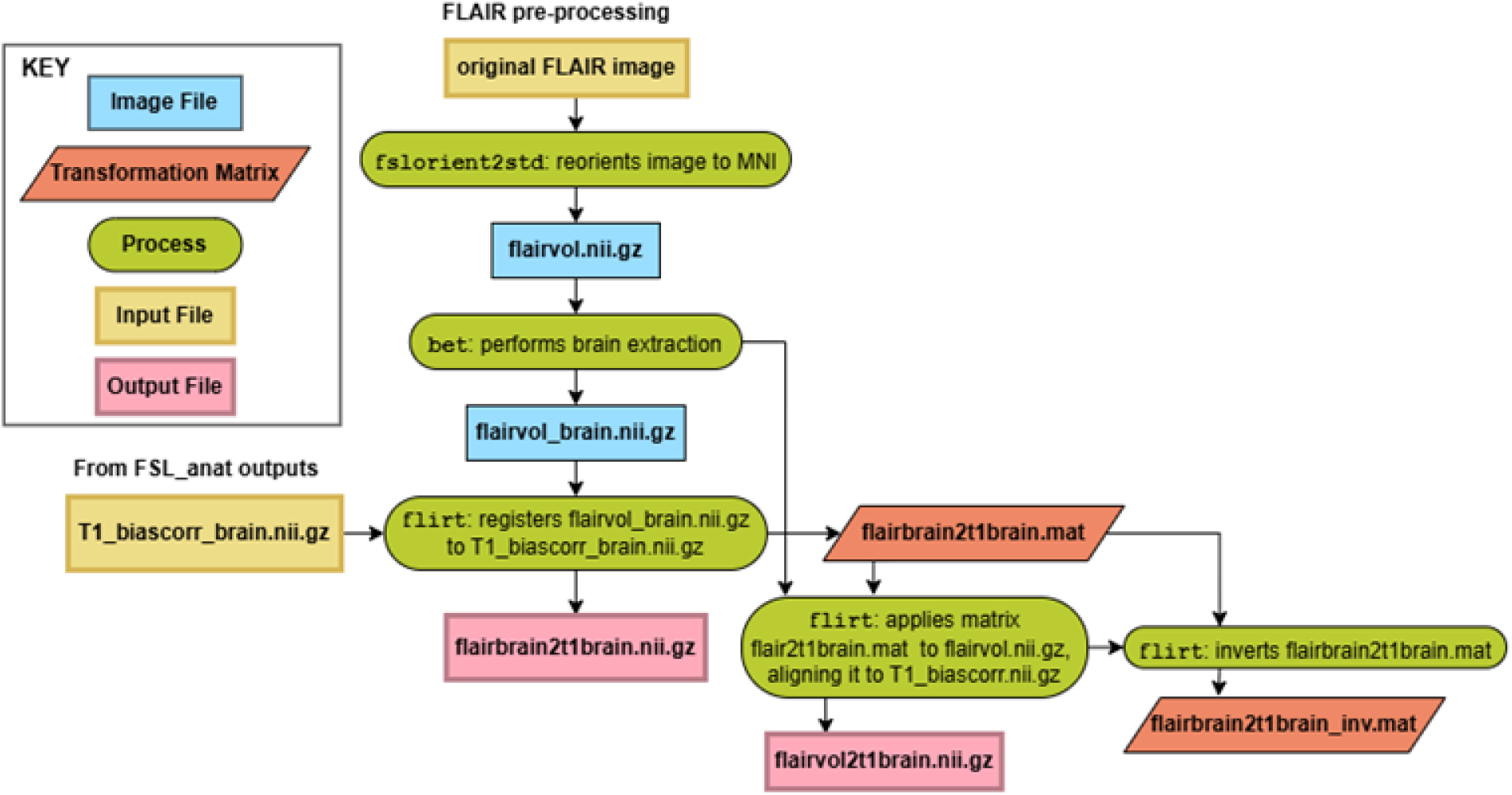
Schematic overview of the preprocessing of the FLAIR images in the ENIGMA-PD-WML pipeline. Blue boxes show image files, red parallelograms show transformation matrices, green ovals show processes, yellow boxes show input files and pink boxes show output files.

**Figure 4:**
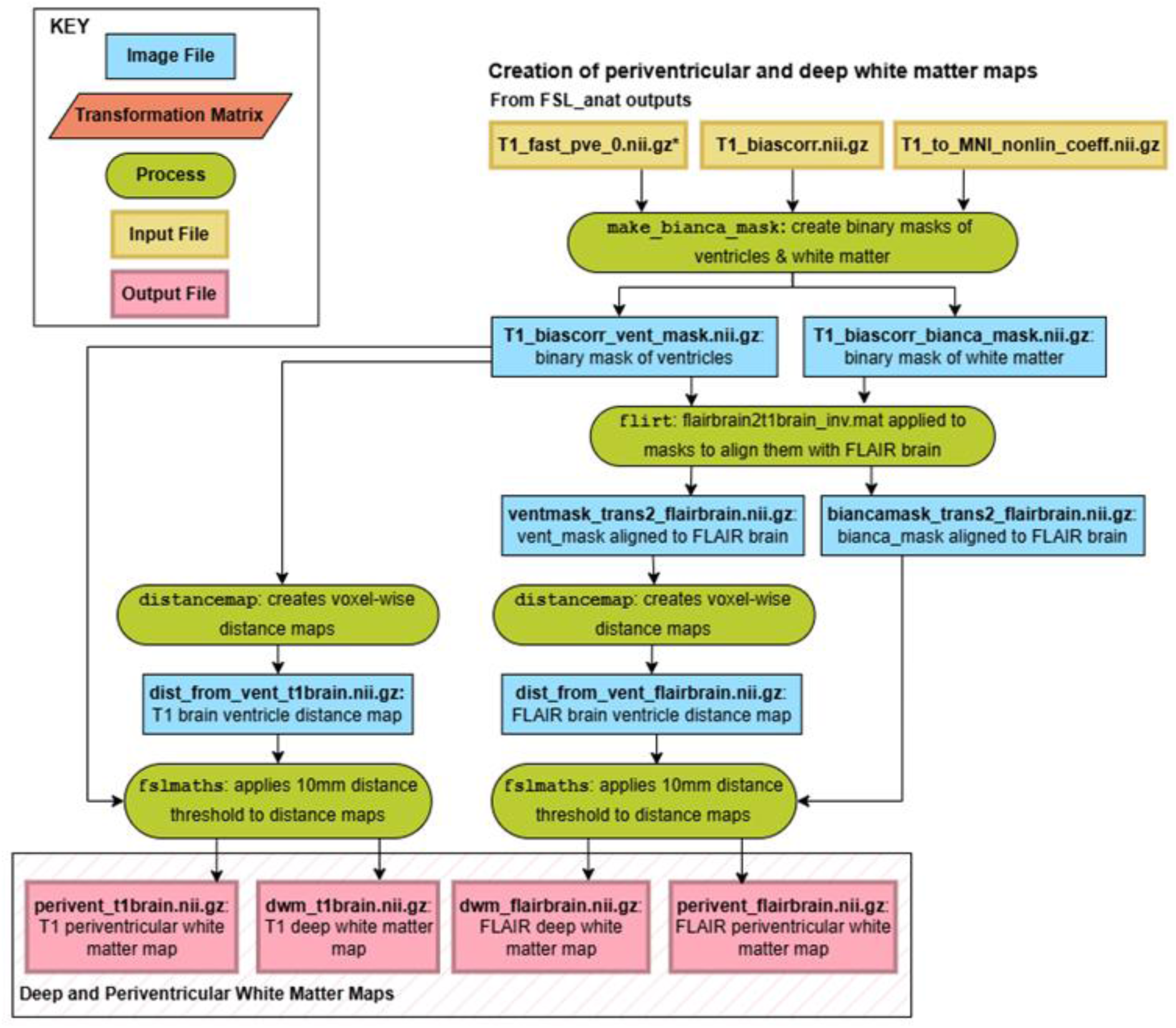
Schematic overview of the steps involved in the creation of deep and periventricular white matter maps in the ENIGMA-PD-WML pipeline. Blue boxes show image files, red parallelograms show transformation matrices, green ovals show processes, yellow boxes show input files and pink boxes show output files.

Pipeline Steps:

1. On the T1-weighted image, fsl_anat is run to reorient the image to match the orientation of the Montreal Neurological Institute standard template *MNI152* (fslorient2std), automatically crop the image (robustfov) and apply radiofrequency and B1-inhomogeneity bias field correction (fast). The image is then both linearly registered (flirt) and then non-linearly warped (fnirt) to MNI space. Following this, brain-extraction (bet) and tissue-type segmentation (fast) is performed.
2. The FLAIR image is also reoriented to MNI orientation (fslreorient2std) and brain extraction performed (bet).
3. The extracted FLAIR brain undergoes 3D rigid body registration to the cropped bias field-corrected extracted T1-weighted brain generated in step 1 using a normalized mutual information cost function and six degrees of freedom (flirt).
4. The transformation matrix generated by this registration is applied to the FLAIR image, aligning it with the cropped T1-weighted image using trilinear interpolation (flirt). The transformation matrix is then inverted for use in step 6.
5. Binary masks of the ventricles, *vent_mask,* and white matter, *bianca_mask*, are created by running make_bianca_mask on the cropped bias field-corrected T1-weighted image, CSF tissue probability map and the MNI-to-cropped T1-weighted non-linear warp field generated from step 1.
6. The inverted FLAIR-to-cropped T1-weighted transformation matrix from step 4 is applied to *vent_mask* and *bianca_mask*, aligning them to the FLAIR image using nearest neighbor interpolation (flirt).
7. Voxel-wise distance maps from the edge of the ventricles are created for both the cropped T1-weighted and original FLAIR images by applying distancemap to each *vent_mask*.
8. Periventricular masks, *perivent*, are created by applying a 10 mm upper threshold to the distance map and deep white matter masks, *dwm*, are created by applying a 10 mm lower threshold to the distance map using fslmaths. *Note: There is an overlap of the periventricular and deep white matter binary masks at 10 mm. The overlapping boundary can be identified by multiplying the masks and subtracting the boundary from the deep white matter mask. This results in the periventricular region defined as 10 mm or less from the edge of the ventricles and the deep white matter region defined as greater than 10 mm from the edge of the ventricles. Large confluent lesions will be divided by this technique into periventricular and deep white matter portions. See ‘Post-Quality Control Step for Large Periventricular Lesions: Clustering’ for details on an optional additional approach for further categorizing these lesions*.
9. To prepare for UNet-pgs, if the cropped T1-weighted image from step 1 has an x- or y-dimension of 500 pixels or more, it and the aligned FLAIR from step 4 are further cropped to 472 x 496 pixels (fslroi). This double-cropped T1-weighted image then undergoes a 3D rigid body registration to the cropped T1-weighted image from step 1 using a normalized mutual information cost function and six degrees of freedom (flirt), to allow for transformations and warping of outputs to MNI space. Pre-processing Rationale: The pre-processing steps have been included in the pipeline to allow incorporation of T1-weighted and FLAIR images with varying acquisition parameters such as field strength, slice thickness, acquisition plane and voxel size. Incorporating these steps is essential to allow the pipeline to be used at various international sites, as MRI acquisition parameters often differ across cohorts. Consequently, a greater number of sites will be able to use the pipeline, particularly on imaging data collected with a different research question in mind, without the need for particular MRI protocols. There is a variation in our implementation of the UNet-pgs approach in that we register the FLAIR image to the T1-weighted rather than *vice versa* (as described in Park et al. (2021)). The rationale is that the MNI template is T1-weighted and our approach cuts a step out by transforming and warping segmentation outputs directly to MNI space. We reorient both the T1-weighted and FLAIR images to match the orientation of the MNI template and also apply a bias field correction to both images to address field inhomogeneities. Additionally, we include a step for the creation of deep and periventricular WML maps, due to the likely clinical significance of the location of WML and potential differences in pathological correlates (Kim et al., 2008) (Kim et al., 2008). We have followed the approach taken by Griffanti et al. (2018) to classify the periventricular and deep white matter based on a 10 mm distance from the lateral ventricles (Griffanti et al., 2018, DeCarli et al., 2005). This classification method is suited to automated segmentation approaches; however, we acknowledge that applying a fixed distance may be limited in the presence of atrophy and ventricular dilation. Finally, all images must be cropped to less than 500×500 pixels before input to UNet-pgs, so that images are small enough for the deep learning model to perform effectively. WML Segmentation using UNet-pgs Following the pre-processing steps outlined above, the pre-processed cropped T1-weighted and aligned FLAIR images are ready for input into the UNet-pgs algorithm (Park et al., 2021) for WML segmentation. A diagrammatic overview of running UNet-pgs within the pipeline is shown in **Figure 5**. Pipeline Steps:
10. UNet-pgs is run on the pre-processed cropped T1-weighted and aligned FLAIR images. The algorithm uses similarity indices and affine transformations and resampling volume to register the images to each other and performs a rough bias field correction. It then calculates the features, classifies the voxels and fits the shape model. The images undergo free deformation before the volume is built from mesh and warped back to its original space.
11. All intermediary outputs from UNet-pgs steps are housed in the input folder. When UNet-pgs is complete, a binary segmentation mask, aligned to the pre-processed cropped T1-weighted image and aligned FLAIR, *results.nii.gz,* is generated for each subject and stored in the output folder. Post-processing Once UNet-pgs is complete, each subject will have a binary WML segmentation mask. A series of post-processing steps using FSL are applied to the WML masks to generate outputs in MNI standard space. **Figure 6** shows the post-processing steps and outputs of the pipeline and **Figure 7** gives an overview of the entire pipeline, including pre-processing, UNet-pgs and post-processing steps. Pipeline Steps:
12. The 1 mm resolution MNI152 standard space T1-weighted average structural template image, *MNI152_T1_1mm*, and extracted brain, *MNI152_T1_1mm_brain*, are copied to the output folder. The 1 mm resolution JHU ICBM-DTI-81 white matter labels atlas, *JHU_ICBM_labels_1mm*, and Oxford-GSK-Imanova striatal probabilistic connectivity atlas, *striatum_con_label_thr50_7sub_1mm*, (segmented into 7 sub-regions and thresholded at 50%) aligned to *MNI152_T1_1mm* are also copied to the output folder. Intermediary files are not automatically deleted, enabling troubleshooting, QC and any further analysis later.
13. The transformation matrices created in steps 1, 6 and 9 are applied to the UNet-pgs outputs to align with the reoriented full field-of-view T1-weighted image, reoriented full field-of-view FLAIR and cropped T1-weighted image respectively using nearest neighbor interpolation (flirt).
14. The UNet-pgs results aligned to the cropped T1-weighted image and reoriented full field-of-view FLAIR from step 12 are split into periventricular and deep white matter results by multiplying with the *perivent* and *dwm* masks from step 8 (fslmaths).
15. The transformation matrix from step 1 is applied to the cropped periventricular and deep white matter results from step 14 to align them with the reoriented full field-of-view T1-weighted image from step 1 using nearest neighbor interpolation (flirt).
16. The T1-to-MNI linear transformation matrix (flirt) and non-linear warp coefficients (applywarp) from step 1 are applied to the cropped whole T1-weighted image, periventricular and deep white matter results from step 14 to align them with *MNI152_T1_1mm* using nearest neighbor interpolation. The T1-to-MNI linear transformation matrix (flirt) and non-linear warp coefficients (applywarp) are also applied to the cropped bias field-corrected extracted brain from step 1 and the aligned FLAIR brain from step 15 to align them with *MNI152_T1_1mm* using trilinear interpolation.
17. The linear periventricular output *results2mni_lin_perivent* is added to the linear deep output *results2mni_lin_deep* from step 16 to create a linear binary combined white matter output *results2mni_lin_combined* (fslmaths). Separately, the non-linear periventricular output *results2mni_nonlin_perivent* is added to the non-linear deep output *results2mni_nonlin_deep* from step 16 to create a non-linear binary combined white matter output *results2mni_nonlin_combined* (fslmaths).
18. The T1-to-MNI non-linear warp coefficients from step 1 are inverted (invwarp) and applied to *JHU_ICBM_labels_1mm* and *striatum_con_label_thr50_7sub_1mm* to align them with the cropped T1-weighted image using nearest neighbor interpolation (applywarp).
19. The results aligned to the cropped T1-weighted image from step 13 are multiplied with the aligned *JHU_ICBM_labels_1mm* and *striatum_con_label_thr50_7sub_1mm* to generate white matter tract and striatal connection partitions, respectively (fslmaths).
20. The T1-to-MNI linear transformation matrix (flirt) and non-linear warp coefficients (applywarp) from step 1 are applied to the cropped white matter tract and striatal connection partitions from step 19 to align them with *MNI152_T1_1mm* using nearest neighbor interpolation.
21. All of the results in MNI space, including the T1-weighted and FLAIR brains aligned with *MNI_T1_1mm,* are compressed into a zip folder.

**Figure 5:**
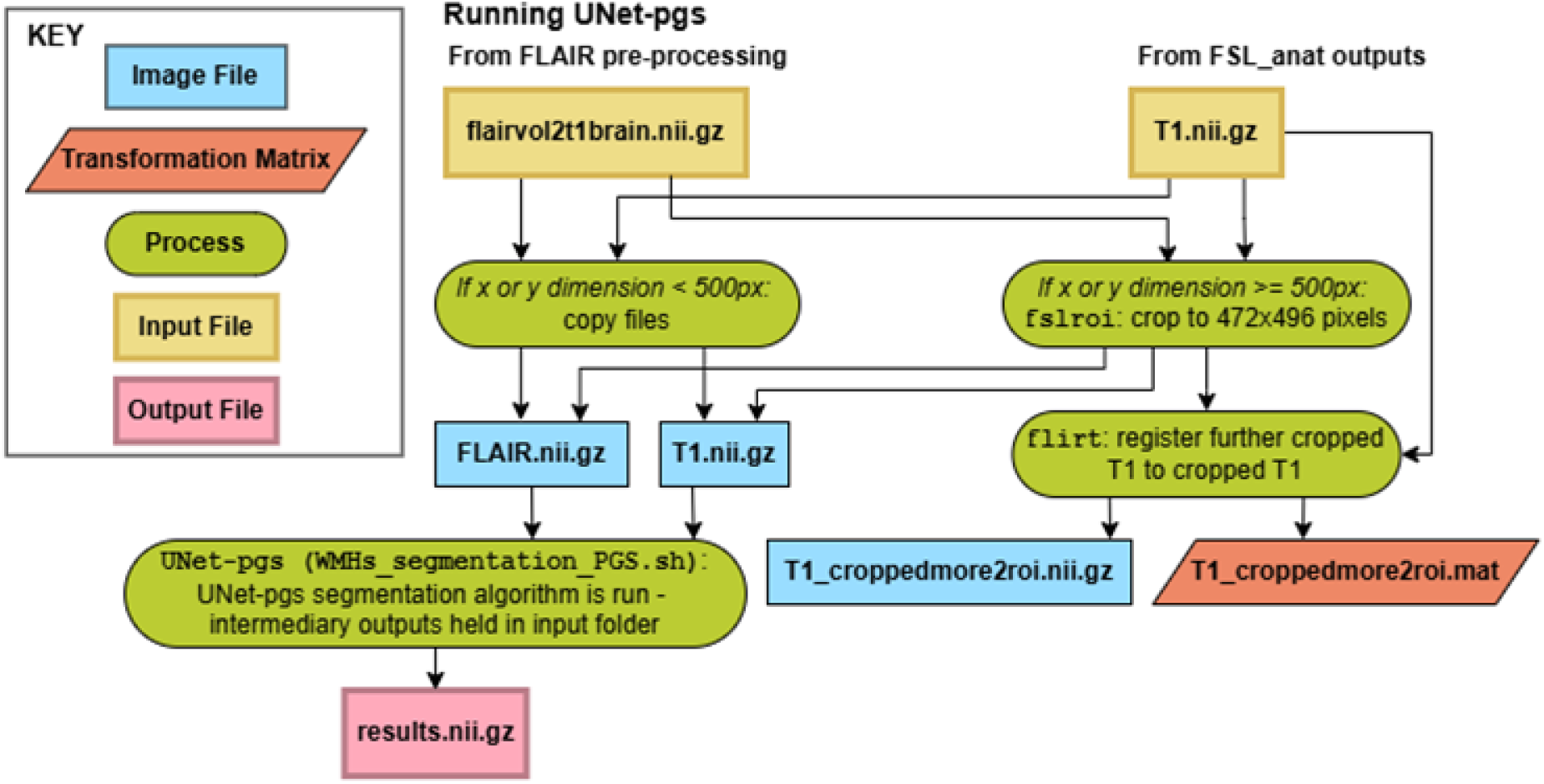
Schematic overview of the steps involved in running the UNet-pgs algorithm within the ENIGMA-PD-WML pipeline. Blue boxes show image files, red parallelograms show transformation matrices, green ovals show processes, yellow boxes show input files and pink boxes show output files.

**Figure 6:**
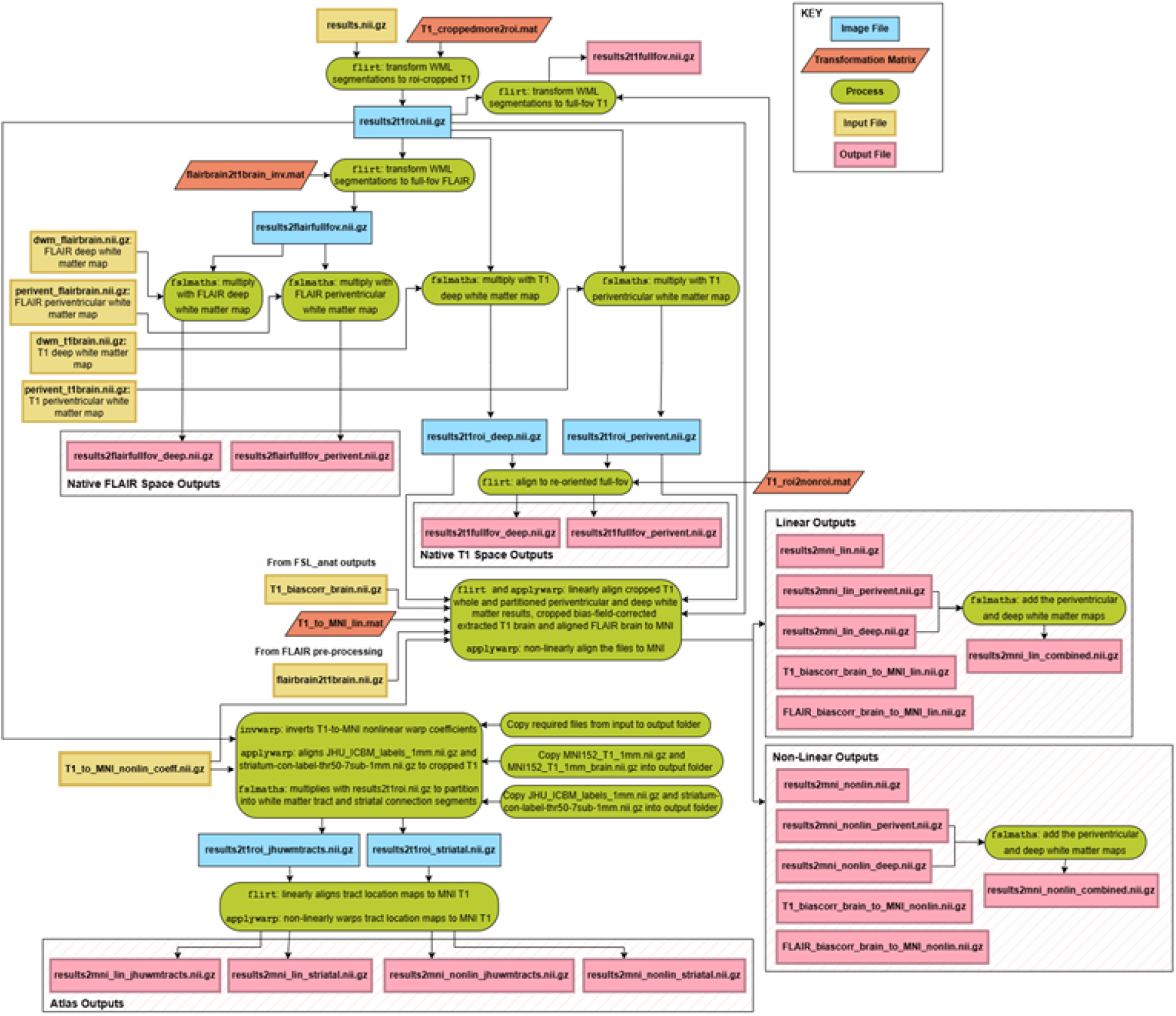
Schematic overview of the post-processing steps of the ENIGMA-PD-WML pipeline. Blue boxes show image files, red parallelograms show transformation matrices, green ovals show processes, yellow boxes show input files and pink boxes show output files.

**Figure 7:**
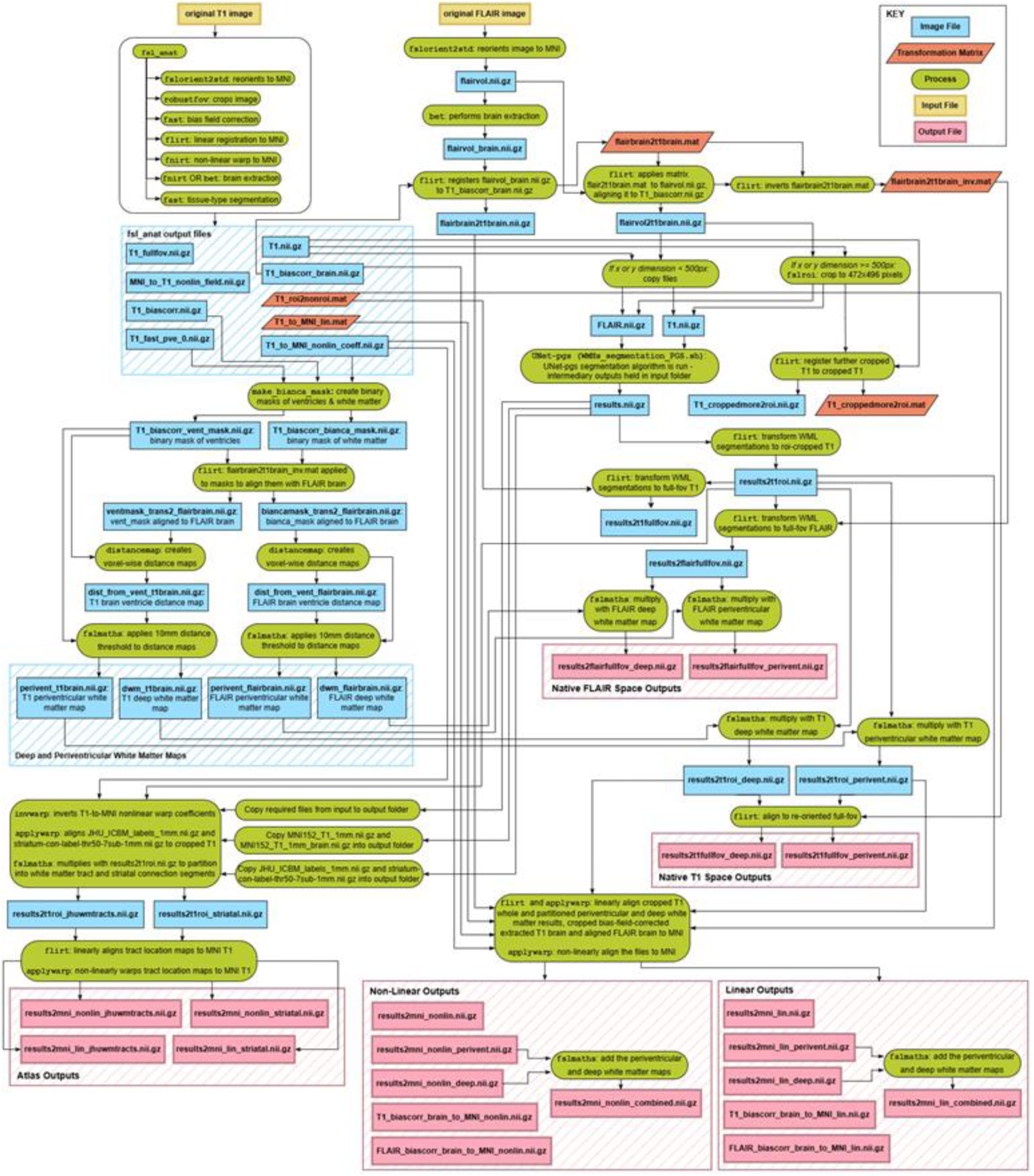
Schematic overview of the entire ENIGMA-PD-WML pipeline. Blue boxes show image files, red parallelograms show transformation matrices, green ovals show processes, yellow boxes show input files and pink boxes show output files.

### Post-processing Rationale

As discussed in the introduction, to allow transfer of data between sites and allow use of the data by consortia, the segmentation outputs are skull and scalp-stripped and transformed to MNI standard space. This ensures the images are anonymized and allows for comparison of neuroanatomical locations across subjects. The images are both linearly transformed to the MNI template for preservation of volumes and non-linearly warped for precise neuroanatomical alignment (Klein et al., 2009). The outputs are also produced in subject-native space for use by the sites where the data originated.

When transforming and warping the binary WML segmentation masks into different spaces, we used nearest-neighbor interpolation. This approach was chosen in order to preserve the binary nature of the segmentation masks. Even so, nearest-neighbor interpolation tends to increase the apparent size of small lesions and potentially shift their location. Alternatively, a trilinear interpolation could be used, but this would require a thresholding step and binarization. The disadvantage of this approach is the potential variability in threshold selection, especially with variable protocols, as would be expected with multi-site data.

We combine the periventricular and deep WML outputs in order to ensure the segmentations are restricted to the white matter and exclude grey matter, CSF and areas outside of the brain. As UNet-pgs works using an intensity approach, this helps to exclude false identification of hyperintensities outside of the white matter as WML.

The inclusion of the John Hopkins ICBM-DTI-81 white-matter labels atlas allows for identification of WML in key white matter tracts in the brainstem as well as in projection, association and commissural fibres (Mori et al., 2005). The Oxford-GSK-Imanova striatal probabilistic connectivity atlas allows for identification of WML in anatomical connections from the striatum to 7 key cortical zones: limbic, executive, rostral-motor, caudal-motor, parietal, occipital and temporal (Tziortzi et al., 2014). Many other atlases also are available in MNI space, allowing users to apply whichever atlas is appropriate without the need for additional transformation.

### Outputs from Pipeline

Upon completion of the pipeline, an enigma-pd-wml folder will be created within the derivatives folder which will be within the root directory. This contains the input and output folders and files for each subject, alongside a log and a zip file containing derived images. An example of these derived outputs in MNI space is shown in **Figure 8**. For each subject the pipeline will produce full WML segmentation maps, deep WML maps and periventricular WML maps. These are all available in both subject-native T1-weighted and FLAIR space and transformed to MNI space both linearly, to preserve volumes, and non-linearly for neuroanatomical alignment.

**Figure 8:**
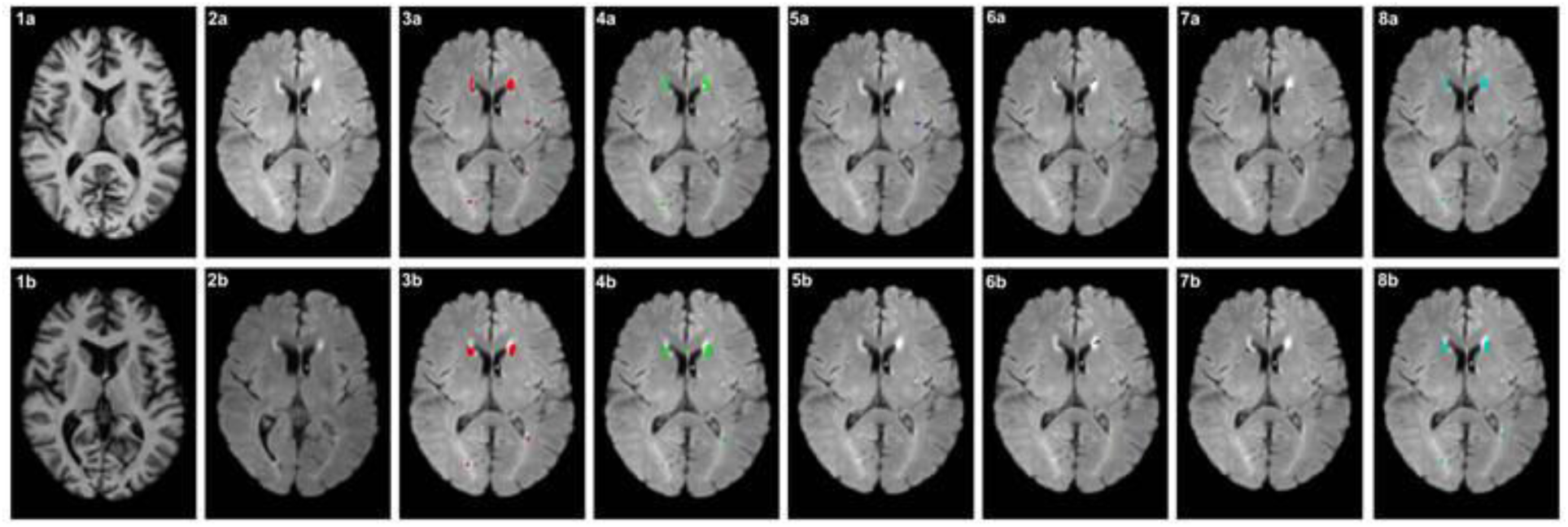
The outputs in MNI space produced by the ENIGMA-PD-WML pipeline for an example subject. The top row images are transformed to MNI space through non-linear warping, the bottom row images through linear registration. 1a: T1 brain non-linearly warped to MNI (*T1_biascorr_brain_to_MNI_nonlin*) 1b: T1 brain linearly registered to MNI (*T1_biascorr_brain_to_MNI_lin*) 2a: FLAIR brain non-linearly warped to MNI (*FLAIR_biascorr_brain_to_MNI_nonlin*) 2b: FLAIR brain linearly registered to MNI (*FLAIR_biascorr_brain_to_MNI_lin*) 3a: Full WML segmentation map non-linearly warped to MNI (*results2mni_nonlin*) 3b: Full WML segmentation map linearly registered to MNI (*results2mni_lin*) 4a: Periventricular WML segmentation map non-linearly warped to MNI (*results2mni_nonlin_perivent*) 4b: Periventricular WML segmentation map linearly registered to MNI (*results2mni_lin_perivent*) 5a: Deep WML segmentation map non-linearly warped to MNI (*results2mni_nonlin_deep*) 5b: Deep WML segmentation map linearly registered to MNI (*results2mni_lin_deep*) 6a: WML in John Hopkins ICBM-DTI-81 white-matter labels atlas non-linearly warped to MNI (*results2mni_nonlin_jhuwmtracts*) 6b: WML in John Hopkins ICBM-DTI-81 white-matter labels atlas linearly registered to MNI (*results2mni_lin_jhuwmtracts*) 7a: WML in Oxford-GSK-Imanova striatal probabilistic connectivity atlas non-linearly warped to MNI (*results2mni_nonlin_striatal*) 7b: WML in Oxford-GSK-Imanova striatal probabilistic connectivity atlas linearly registered to MNI (*results2mni_lin_striatal*) 8a: Combined periventricular and deep WML segmentation map non-linearly warped to MNI (*results2mni_nonlin_combined*) 8b: Combined periventricular and deep WML segmentation map linearly registered to MNI (*results2mni_lin_combined*)

In addition to the image outputs above, the pipeline also outputs a CSV file (*data/derivatives/enigma-pd-wml/t1_volumes.csv*) containing the estimated brain volume (in mm³) in native T1-weighted space and MNI space, as well as the factor used to scale the brain from T1-weighted to MNI space for each subject and session. These values are generated from fsl_anat and can be used for head-size normalization in longitudinal analyses.

## Quality Control

As with any automated pipeline, some form of manual quality control (QC) is required to ensure the pipeline has performed correctly and the outputs are adequate for further analysis. The original QC process for the pipeline consisted of an instruction document outlining how to open the FLAIR images and overlay the WML segmentation maps to assess the quality of the segmentation, with image examples of good and poor segmentations. However, this approach, which involved manually opening images for each subject in MRI viewing platforms, required substantial time and effort.

To make the QC procedure more efficient and straightforward, an interactive QC process was developed and embedded in the pipeline. Firstly, 2D PNG images are generated from the 3D NIfTI images, then assembled into an interactive browser-based HTML interface to allow for fast and easy visualization.

Opening of the HTML allows for visual examination of the WML segmentations by overlaying the WML masks (*results2mni_nonlin_combined.nii.gz* and *results2mni_lin_combined.nii.gz*) onto the FLAIR images registered to MNI space (*FLAIR_biascorr_brain_to_MNI_nonlin.nii.gz* and *FLAIR_biascorr_brain_to_MNI_lin.nii.gz*) [**Figure 9**]. Additionally, the HTML allows for review of both linearly registered and non-linearly warped outputs separately, to assess the quality of the transformation. The interface incorporates built-in tools allowing for navigation through 182 slices, toggling ON/OFF of the overlay and QC rating. All images can be marked as a PASS, FAIL or FLAG FOR LATER QC. Users can select failure reasons from the dropdown box or provide additional comments using the free-text box. A QC guide is available within the interface which provides example images for each of the failure categories, alongside examples of good segmentations to ensure standardized failure categorization across sites. The interface loads up to 200 subjects and has an auto-save function which saves the page every 30 seconds. Following completion, a timestamped CSV file can be downloaded with the QC decisions. Full technical details and a usage guide are available at https://github.com/UCL/Enigma-PD-WML/blob/main/docs/qc_pipeline.md and https://github.com/UCL/Enigma-PD-WML/blob/main/docs/qc_usage.md, respectively.

**Figure 9:**
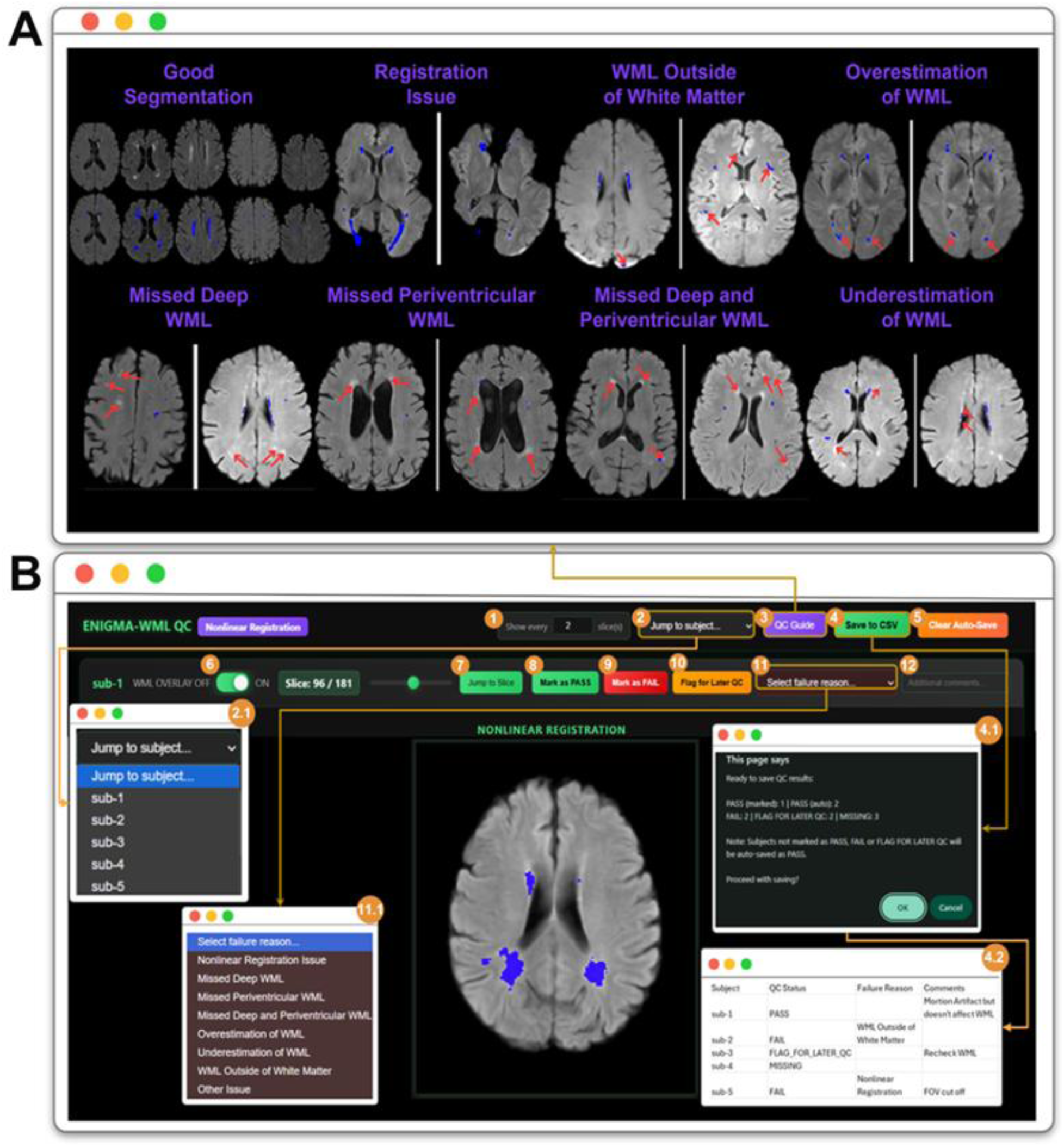
ENIGMA-PD-WML QC HTML Interface and QC Guide. *Panel A:* The QC Guide reference panel displays example images for each QC category. Categories include: Good Segmentation (acceptable WML segmentation), Registration Issue (misalignment between FLAIR and MNI space), WML Outside of White Matter (WML segmentations in gray matter or CSF), Overestimation of WML (segmentation extends beyond true lesion boundaries), Missed Deep WML (undetected lesions in deep white matter), Missed Periventricular WML (undetected lesions adjacent to ventricles), Missed Deep and Periventricular WML (both regions affected), and Underestimation of WML (incomplete lesion coverage). Blue overlay indicates WML segmentation; red arrows highlight areas of concern. *Panel B*: The QC interface with feature controls: (1) Slice skip interval selector (2) Subject dropdown for navigating to a specific subject of interest (2.1) Subject selection dropdown listing available subjects (3) QC Guide toggle button, always available for review (4) Export to CSV button (4.1) Save to CSV confirmation dialog box summarizing QC decisions (PASS, FAIL, FLAG FOR LATER QC, and MISSING counts) before CSV export. (4.2) Exported CSV format containing Subject ID, QC Status, Failure Reason, and Comments columns. (5) Clear Auto-Save button, in case the user wants to erase previously saved data; (6) WML overlay ON/OFF toggle; (7) Slice navigation slider (displaying slice 96 of 181, starts from 0) along with Jump to specific slice; (8) Mark as PASS button; (9) Mark as FAIL button; (10) Flag for Later QC button; (11) Failure reason dropdown (required when marking FAIL). (11.1) Failure reason dropdown showing all categories. (12) Free-text box for additional notes.

### Optional Post-Pipeline Analysis Approaches

#### Post-Quality Control Step for Large Periventricular Lesions: Clustering

As mentioned in the pre-processing steps, the pipeline itself uses distance mapping to generate periventricular and deep WML maps. This approach classifies any lesions less than 10 mm from the ventricles as periventricular, and all others as deep. This is a widely adopted approach for automated segmentation tools (Griffanti et al., 2018); however, when inspecting outputs we found that large confluent lesions were arbitrarily split at this distance, resulting in a singular lesion being classified as both periventricular and deep. To overcome this, we developed a clustering method for use following QC to correctly categorize these lesions as confluent.

This method combines the fixed distance approach with a new approach that labels confluent lesions as distinct components within a binary lesion mask to ensure continuous lesions are classified under a singular identity. Using the confluent lesion method, ventricle segmentations are dilated at a fixed distance of 10 mm in all directions, accounting for anisotropic voxel resolution. Lesions that are split at this distance are then labeled confluent, resulting in a third output map: confluent. This is included as a separate module which can be applied following QC to produce confluent outputs in subjects with a total lesion volume over 10 cm^3^.

The lesion clustering and classification module is implemented in Python (version ≥ 3.8) and can be executed from the command line, following completion of the pipeline. The script requires installation of the following dependencies: NumPy, SciPy, NiBabel, and scikit-image. Execution of the clustering module performs batch processing across all subjects defined in the input directory structure.

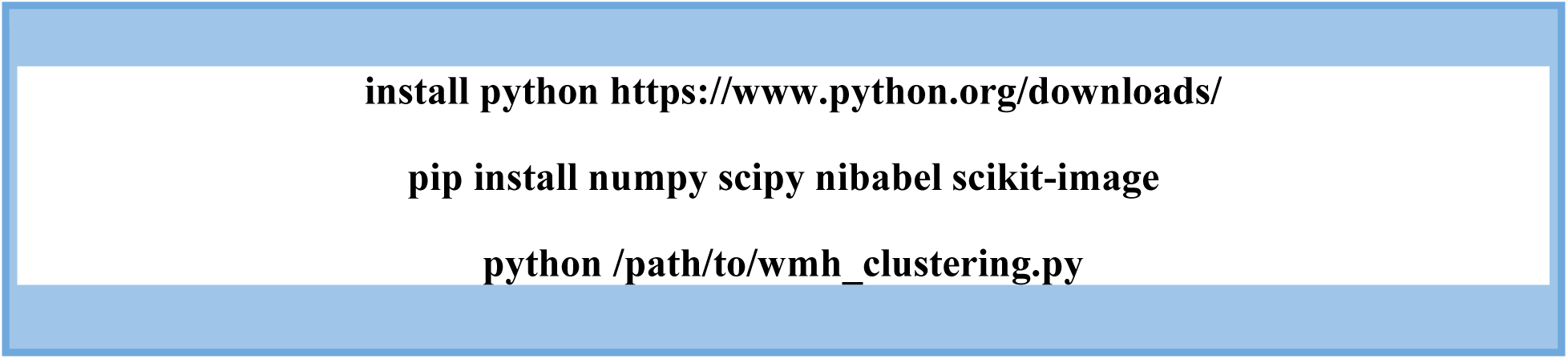

The code for this module is available alongside the main pipeline on GitHub (https://github.com/UCL/Enigma-PD-WML#periventricular-and-deep-wml-clustering).

#### Visualization of WML Distribution: Bullseye

As discussed in the introduction, lesion location is now recognized to be clinically important (Meng et al., 2022). Therefore, the outputs from the pipeline identify lesions in the periventricular region and deep white matter, as well as in white matter tracts. Although not part of the pipeline, we have looked into methods that can be applied to WML segmentation outputs from the pipeline to visualize the distribution of WML. A potential method is use of the Bullseye tool, developed by Sudre et al. (2019). Bullseye parcellates the brain into four equidistant layers and nine lobar zones resulting in 36 regions in which WML burden can be assessed. The results can be displayed in a Bullseye plot [**Figure 10**] which visually represents the regional distribution of WML. This approach was initially developed using each subject’s T1-weighted image to create patient-specific frames. We tested an adapted Bullseye approach, which created the 36 regions using the T1-weighted MNI template, thereby allowing Bullseye to be applied to the outputs in MNI space. On a sample of 79 subjects (45 PD, 34 healthy controls) Spearman’s correlation analyses found that lesion volumes generated from applying Bullseye to outputs in MNI space correlated strongly with the results of applying Bullseye to outputs in native T1 space (ρ = 0.994, 95% CI 0.98-1.00). Bland-Altman plots also showed agreement between approaches (median bias = 0.0001, limits of agreement −0.0004 to 0.0023). We did, however, see a ‘funnel’ effect in the plots, whereby as the median total WML volume increased, so did disagreement between methods. There was a general underestimation of WML volume in the MNI approach compared to T1, likely due to the nearest-neighbor interpolation approach used to transform the maps in subject-native to MNI space. However, no lesions were missed entirely, the limits of agreement contained zero, and there was a strong monotonic linear relationship between approaches. As such, we believe bullseye can be used in MNI space whilst acknowledging these limitations and the potential reasons for these differences.

**Figure 10:**
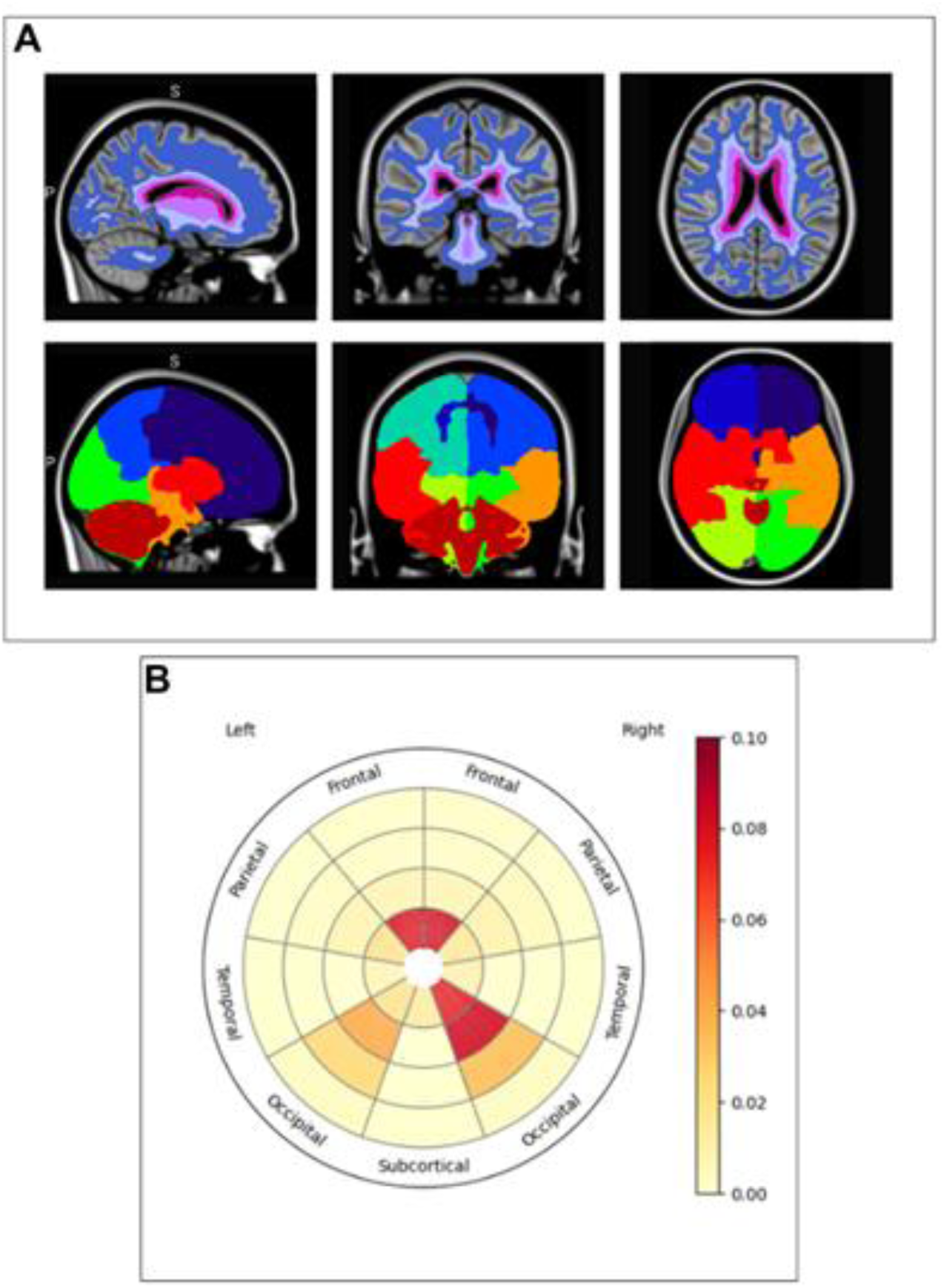
Bullseye segmentation and visual representation. *Panel A*: Bullseye parcellation in MNI space. The top row displays the four layers from the ventricles to the cortical grey matter. The bottom row displays the nine lobar regions: left frontal, right frontal, left parietal, right parietal, left occipital, right occipital, left temporal, right temporal and combined basal ganglia, thalami and infratentorial region (BGIT). *Panel B:* Bullseye plot, displaying the median lesion frequency per region. More red regions represent greater frequency; more yellow regions represent lower frequency. The layer closest to the center of the plot represents the most periventricular layer; the outer layer represents the layer which meets the cortex, totaling 36 regions moving the periventricular area to the cortex.

## Discussion

The ENIGMA-PD-WML containerized pipeline is a streamlined, standardized approach to create reproducible WML maps. The pipeline requires minimal manual input aside from converting images to NIfTI format and structuring the data for input to the pipeline. All relevant code and software are within the containerized package, allowing the pipeline to run using a singular line of code from the command line. This approach minimizes the need for technical expertise and extensive software downloads, allowing the pipeline to be ‘pluggable’ and therefore suitable for use in various settings. There are also options to modify the default running of the pipeline, by running subjects in parallel or with the addition of optional flags. The pipeline is freely available for use at https://github.com/UCL/Enigma-PD-WML.

The only neuroimaging requirements are a T1-weighted and FLAIR image for each subject with a slice thickness of 5 mm or less, taken from an MR scanner with field strength of at least 1.5T to ensure accurate WML detection (Duering et al., 2023, Wardlaw et al., 2013). Various steps ensure the pipeline works on any T1-weighted and FLAIR images that meet these basic requirements, thereby accounting for varying scanner parameters and protocols. This is essential to allow data from multiple research groups to be collated and assessed using a standardized methodological approach. Furthermore, the pipeline produces all outputs in both subject-native and standard MNI space. These outputs include full segmentation maps as well as periventricular and deep white matter maps and maps of WML located within regions of the John Hopkins ICBM-DTI-81 white-matter labels atlas and the Oxford-GSK-Imanova striatal probabilistic connectivity atlas to aid lesion localization. MNI images can be easily shared as anonymized data, allowing for group comparisons, as well as identification of individual differences between subjects. This ensures the pipeline is ideal for large-scale data analysis, in which data is shared from various sites, as well as for site-specific use.

**Figure 7** visually illustrates the extensive additional steps performed by the code in order for the user to simply structure and input the data without the need for any further manual pre-processing or post-processing steps. In brief, the pre-processing steps performed by the code register the T1-weighted and FLAIR image together and crop these to less than 500 pixels. Both of these steps are required for running the UNet-pgs algorithm and would otherwise need to be done manually. Transformations for registration of the images to MNI space and white matter atlases are then generated. We use an adapted version of UNet-pgs in which we register the FLAIR to the T1-weighted image, rather than registering the T1-weighted image to the FLAIR (Kuijf et al., 2019, Park et al., 2021). Despite WML being most visible in FLAIR acquisitions, registering to the T1-weighted image helps to preserve anatomical detail, as often the T1-weighted images have better resolution and contrast than FLAIR images, and prepare for registration to MNI as the MNI template is T1-weighted.

From a pragmatic perspective, development of this pipeline required a multi-disciplinary approach including clinicians, image analysis specialists, software engineers and consortia leads. The title of the pipeline reflects the ENIGMA-PD working group to which the team that developed the pipeline belong, yet we expect that any group interested in WML of presumed vascular origin would benefit from the pipeline as it is based on UNet-pgs with evidence for its successful segmentations in multiple disease groups (Kim et al., 2024, Joo et al., 2022, Park et al., 2021). Whether the pipeline is suitable for neuroinflammatory or other WML etiologies is yet to be determined.

### Our Experience of Using the ENIGMA-PD-WML Pipeline

To date, the pipeline has been applied to over 2000 subjects from 8 different cohorts within the ENIGMA-PD consortium. We completed preliminary analyses on over 400 subjects with Parkinson’s disease and 250 healthy controls from 6 cohorts (Al-Bachari et al., 2026a). Most sites, regardless of clinical or imaging expertise, were able to implement the pipeline with minimal additional support. Depending on site-specific preferences we arranged an initial meeting to help guide them through the process; a few sites required a follow-up meeting to troubleshoot and discuss errors. All sites that applied the pipeline were able to share data from the majority of their datasets. The general experiences were that the pipeline was straightforward to run and performed well. However, as with any automated neuroimaging approach, there was a small subset of scans which produced errors, as outlined below. It is important to note the QC process is an integral part of the pipeline, ensuring any incorrect segmentations are clearly identified and excluded from any further analysis.

If the original images have poor contrast between the white matter and grey matter, this can cause UNet-pgs to misidentify parts of the cortex as WML. To overcome this issue, we apply a white matter mask to the outputs in order to restrict the segmentations to the white matter only in the combined periventricular and deep white matter lesion output. However, this is reliant on the contrast of the original image being good enough to obtain an accurate white matter mask.

Issues with non-linear warping to MNI standard space have been noted when FSL’s brain extraction tool has not extracted the full brain adequately from either the original T1 or FLAIR image, which in turn can cause uninterpretable outputs from UNet-pgs. The current brain extraction process has been optimized to work in a range of scanner parameters; however, inevitably there will be some exceptions.

On rare occasions, UNet-pgs generates unexpected WML segmentations, such as missing visually obvious WML, misidentifying hyperintensities outside of the white matter on the FLAIR images as WML, and overestimation of the optic radiation as WML. Another observation we have made is that when the original images have been acquired with a tilt in the imaging plane this can cause the outputs from the pipeline to be inaccurate. This may be due to UNet-pgs being trained on data that does not include these examples. After discussion with the creators of UNet-pgs, we jointly feel a potential solution is to generate correct manual segmentations of these cases that can then be used for retraining and fine-tuning of the model.

### Limitations and Common Challenges

The most significant challenge arising from the pipeline is that it is a memory-intensive process, requiring a high-end processor with a minimum of 32GB of RAM and taking approximately 30 minutes per instance, excluding QC. This can generally be overcome by using HPC systems where possible, allowing a substantial amount of RAM to be allocated to the process, without depending on an individual’s computer specifications. Additionally, multiple cores or nodes can be used if available. The pipeline can be run in parallel to enable multiple instances to be processed at the same time. This speeds up overall analysis time but also increases the amount of memory required. Alternatively, for those without access to HPC systems, as the pipeline is fully containerized, it is portable and can be run on a subset of subjects on multiple computers.

Intermediary files are not automatically deleted by the pipeline to enable troubleshooting, QC and any further analyses at a later stage. The input, intermediary and output files take up approximately 800MB of storage per subject per session, assuming all image files are in gzipped NIfTI format. Sites are of course free to delete these files manually as soon as they are no longer needed.

We chose to apply bias field inhomogeneity correction to both the T1-weighted and FLAIR images as an initial pre-processing step. Applying such a correction can sometimes mask WML, especially when large lesions are present, such as infarcts (Valdés Hernández et al., 2016). However, we opted to apply this correction due to the nature of this pipeline intended to be used across sites and imaging protocols.

Another important consideration when assessing the outputs in MNI space, is that non-linear warping from the subject-native to MNI space may affect WML volumes. This is an acknowledged limitation of fitting T1-weighted/FLAIR images to the MNI template, as compression or stretching of lesions may cause slight alterations in volume. These volume changes may also result from using nearest-neighbor interpolation; however, this is required to maintain the binary values of the segmentation (Mahmoudzadeh and Kashou, 2013). Non-linear warping is essential for maintaining precise neuroanatomical location of lesions, which is increasingly considered more relevant to clinical outcomes than volume. However, to ensure analyses of both location and volume can be performed, outputs linearly aligned to MNI are also output from the pipeline and can be used for precise volume estimates.

## Conclusion

The ENIGMA-PD-WML pipeline generates binary WML maps in a standardized way, which can be used for assessment of WML burden and location. The pipeline has been tested on varying scanner parameters and disease populations in the context of WML of presumed vascular origin and allows for sharing of derived data for multi-center imaging analyses, therefore we anticipate its use in several settings. Accurately identifying WML pathology and understanding associations with clinical features will help to unlock the vascular contribution to disease processes. As vascular factors are potentially modifiable, WML can be an effective tool in pathophysiological and therapeutic studies alike. We envisage the ENIGMA-PD-WML pipeline will be a useful resource for both our research and clinical colleagues.

## Acknowledgements

Special thanks to the Academy of Medical Sciences for directly supporting the project. With thanks to Professor Laura Parkes and all members of the ENIGMA-PD core team for their ongoing support. Many thanks also to the ENIGMA-PD sites who have provided invaluable insights into the application of the pipeline.

The authors acknowledge the use of the UCL Myriad High Performance Computing Facility (Myriad@UCL), and associated support services, in the completion of this work and the assistance given by Research IT and the use of the Computational Shared Facility at The University of Manchester. With special thanks to the Centre for Advanced Research Computing at UCL.

## Data Availability Statement

This toolbox paper does not rely on any data. Code and documentation are available at https://github.com/UCL/Enigma-PD-WML.

